# Integrin activation is an essential component of SARS-CoV-2 infection

**DOI:** 10.1101/2021.07.20.453118

**Authors:** Peter Simons, Derek A. Rinaldi, Virginie Bondu, Alison M. Kell, Steven Bradfute, Diane Lidke, Tione Buranda

**Affiliations:** Department of Pathology, University of New Mexico School of Medicine, Albuquerque, New Mexico, 87131, USA; Molecular Genetics and Microbiology, University of New Mexico School of Medicine, Albuquerque, New Mexico, 87131, USA; Department of Internal Medicine, University of New Mexico School of Medicine, Albuquerque, New Mexico, 87131, USA; Center for Infectious Diseases and Immunity, University of New Mexico School of Medicine, Albuquerque, New Mexico, 87131, USA; Comprehensive Cancer Center, University of New Mexico Health Sciences Center, Albuquerque, NM 87131, USA

## Abstract

Cellular entry of coronaviruses depends on binding of the viral spike (S) protein to a specific cellular receptor, the angiotensin-converting enzyme 2 (ACE2). Furthermore, the viral spike protein expresses an RGD motif, suggesting that cell surface integrins may be attachment co-receptors. However, using infectious SARS-CoV-2 requires a biosafety level 3 laboratory (BSL-3), which limits the techniques that can be used to study the mechanism of cell entry. Here, we UV-inactivated SARS-CoV-2 and fluorescently labeled the envelope membrane with octadecyl rhodamine B (R18) to explore the role of integrin activation in mediating both cell entry and productive infection. We used flow cytometry and confocal fluorescence microscopy to show that fluorescently labeled SARS-CoV-2^R18^ particles engage basal-state integrins. Furthermore, we demonstrate that Mn^2+^, which activates integrins and induces integrin extension, enhances cell binding and entry of SARS-CoV-2^R18^ in proportion to the fraction of integrins activated. We also show that one class of integrin antagonist, which binds to the αI MIDAS site and stabilizes the inactive, closed conformation, selectively inhibits the engagement of SARS-CoV-2^R18^ with basal state integrins, but is ineffective against Mn^2+^-activated integrins. At the same time, RGD-integrin antagonists inhibited SARS-CoV-2^R18^ binding regardless of integrin activity state. Integrins transmit signals bidirectionally: ‘inside-out’ signaling primes the ligand binding function of integrins via a talin dependent mechanism and ‘outside-in’ signaling occurs downstream of integrin binding to macromolecular ligands. Outside-in signaling is mediated by Gα_13_ and induces cell spreading, retraction, migration, and proliferation. Using cell-permeable peptide inhibitors of talin, and Gα_13_ binding to the cytoplasmic tail of an integrin’s β subunit, we further demonstrate that talin-mediated signaling is essential for productive infection by SARS-CoV-2.

## INTRODUCTION

Severe acute respiratory syndrome coronavirus 2 (SARS-CoV-2) is a novel virus in the *Betacoronavirus* genus that causes coronavirus disease 2019 (COVID-19).^1^ SARS-CoV-2 was first reported in Wuhan, China, and currently persists as a global pandemic.^2, 3^ SARS-CoV-2 presents similar characteristics with the original SARS-CoV in genome structure, tissue tropism, and viral pathogenesis. However, SARS-CoV-2 is more transmissible than SARS-CoV.

Cellular entry of coronaviruses depends on binding of the viral spike (S) protein to a specific cellular receptor, the angiotensin-converting enzyme 2 (ACE2)^4, 5^ and subsequent S protein priming by cellular protease activity such as Transmembrane Serine Protease 2 (TMPRSS2).^6^ Interestingly, ACE2 expression across different human tissues^7^ revealed low expression of ACE2 in the lungs compared to elevated expression in the kidney and heart.^8, 9^ Nevertheless, studies have shown that type I and II interferons (IFNs) secreted during viral infection upregulate the transcription and expression of ACE2.^10, 11^ Unlike its predecessor, SARS-Cov-2 expresses a novel K403R spike protein substitution encoding an Arginine-Glycine-Aspartic acid (RGD) motif,^4^ introducing the potential for interacting with RGD-binding integrins, as likely mediators for viral cell entry and enhanced pathogenicity.^12^ ACE2 contains two integrin-binding domains: an RGD at position 204–206 and the sequence RKKKNKAR in the cytoplasmic tail at its C-terminus.^13^ Also, ACE2 binds integrin β_1_ in the failing human heart.^13^ Correlated increased expressions of β_1_ ^14^ and ACE2 have been reported.^15, 16^ Others have shown that ACE2 interacts in *cis* with integrin β_1_ in a manner that enhances RGD-mediated cell adhesion.^17^

Integrins are heterodimeric transmembrane adhesion protein receptors composed of α and β subunits whose activation is tightly regulated and bidirectional.^18^ Integrins can exist in three states characterized by their structural conformation and affinity for their ligands (**Fig 1A**). The inactive, bent-closed state (BCS), with a closed headpiece has low affinity for extracellular matrix (ECM) ligands. The bent structure inhibits the receptors from inappropriate signaling due to random binding to extracellular matrix proteins. When primed, integrins exhibit an extended-closed state (ECS) with a closed headpiece and higher ligand binding affinity than BCS. Active and extended-open state (EOS) presents an open headpiece and maximum affinity for ECM ligands.^19^ Integrin function involves coordination with cytoskeletal components whose functions regulate cell adhesion and migration.^20, 21^ Changes in integrin conformation can elicit cell-signaling events that increase ligand affinity/avidity, promote cytoskeletal rearrangement and enable virus internalization. Ligand binding to integrins is mediated by divalent-cations bound at the Metal Ion Dependent Adhesion Site (MIDAS) domain on top of either the αI domain, in I domain-containing integrins, or the βI domain in non-αI integrins.^22^ Physiologically, 1 mM Ca^2+^ and 1 mM Mg^2+^ in body fluid stabilize the BCS conformation. Under non-physiological conditions, 1 mM Mn^2+^ initiates and stabilizes ECS conformation even in the presence of Ca^2+^.

**Figure 1.**
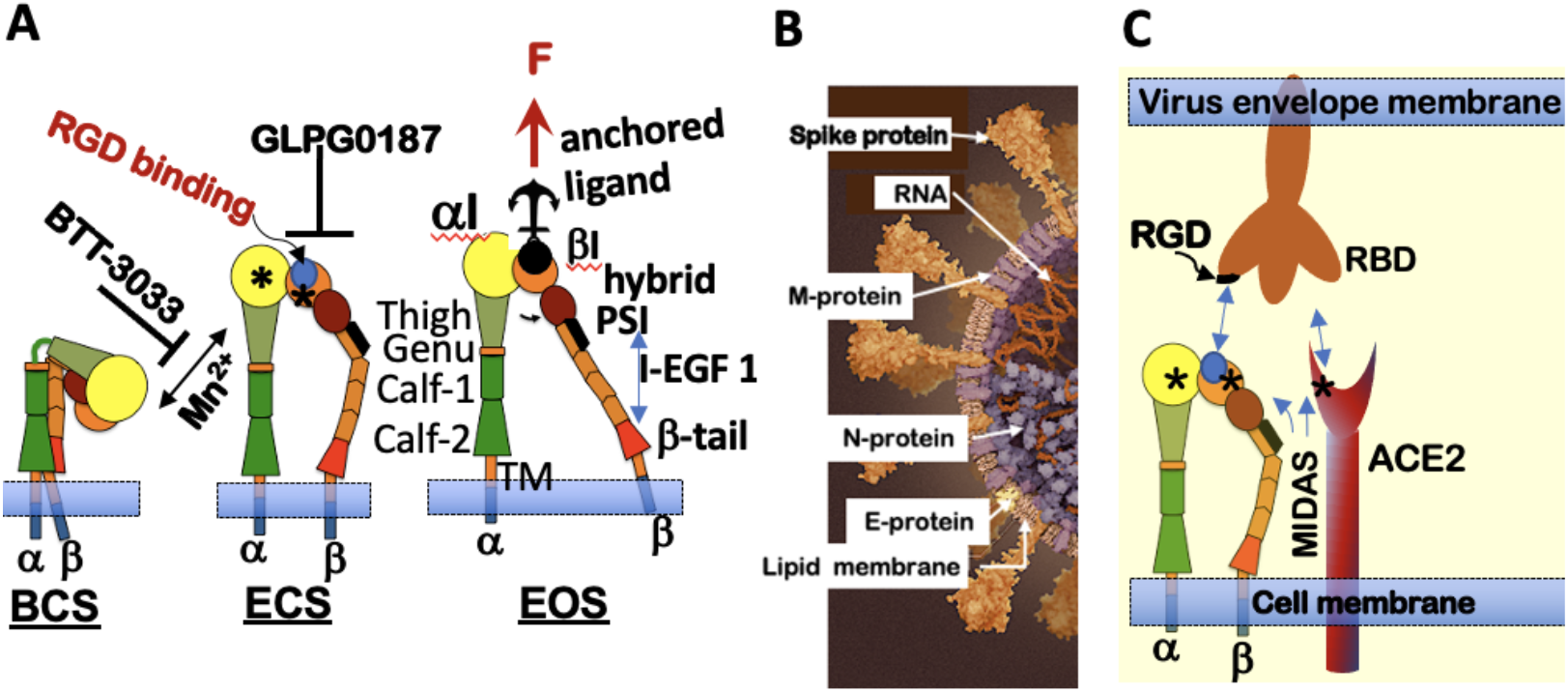
Integrin conformational states antagonist targets and SARS-CoV-2 binding. **A.** *<u>Integrin States</u>*: First, the inactive, bent-closed state (BCS), with a closed headpiece and low affinity for extracellular matrix (ECM) ligands. The bent structure inhibits the receptors from inappropriate signaling due to random binding to extracellular matrix proteins. In the BCS form, binding to large ligands is likely limited. Second, when primed, integrins exhibit an extended-closed state (ECS) with a closed headpiece and higher ligand binding affinity than BCS. Third, active and extended-open state (EOS) with an open headpiece and maximum affinity for ECM ligands. *Integrin Affinity Regulation:* Mn^2+^ binding to the MIDAS site at the αI and βI domain integrin induces integrin extension. α_2_β_1_ integrin antagonist BTT 3033 binds to the a-I domain, and stabilizes the BCS. GLP0187 blocks binding to the RGD ligandbinding domain. EOS binding to a macromolecular ligand or ECM generates force (F) transmitted through the integrin β subunit. **B.** Model of Sars-CoV-2 virion structure (https://www.scientificamerican.com/interactive/inside-the-coronavirus/). SARS-CoV-2 are spherical or ovoid particles of sizes that span the range of 60-140 nm. The SARS-CoV-2 virion consists of a lipid bilayer envelope membrane covering a large nucleoprotein (N)-encapsidated, positive-sense RNA genome. The lipid envelope is decorated with three transmembrane proteins consisting of trimeric spike proteins (S) that project above the lipid bilayer membrane and relatively small membrane (M) and envelope (E) proteins.^64, 65^ S proteins bind with high-affinity (1-50 nM)^4^ to the angiotensin-converting enzyme 2 (ACE2) for productive infection.^66^ **C.** Cartoon alignment of the receptor-binding domain (RBD) and RGD sequence on the trimeric spike protein, which favors engagement of activated integrin, adapted from *ref*. ^24^

Many viruses use integrin-mediated endocytosis pathways for cell entry.^5, 23^ A recent bioinformatics-driven study predicted a model that placed integrins in a central ligating role, whereby SARS-CoV-2 could engage multiple receptors and form a multicomponent receptor complex and functional signaling platform.^24^ Interestingly, ACE2 also has a similar MIDAS motif (**Fig. 1B-C**) ^24^ but it has not yet been established whether the ACE2 MIDAS domain has a potential role in creating synergy overlap between the ligand-binding profiles and regulation of ACE2 and integrins. ^24^ Several *in vitro* studies have established experimental evidence in support of cognate binding interactions between SARS-CoV-2 spike proteins, integrin β_1_ ^25, 26^ and integrin β_3_. ^27, 28^ In addition, the transmembrane glycoprotein neuropilin 1 (NRP1), which is abundantly expressed in the olfactory epithelium and promotes the endocytosis of activated α_5_β_1_ integrin ^29-34^, has been recently identified as a receptor for SARS-CoV-2 infection. ^34, 35^

In this study, we took a mechanistic approach to examine the role of integrins as effectors of SARS-CoV-2 cell entry and productive infection. First, we tested whether inducing a BCS to ECS integrin conformational change with Mn^2+^ ^23, 36^ enhanced cell binding and entry of fluorescently tagged UV-inactivated SARS-CoV-2^R18^. Conversely, we used integrin extension or RGD-binding inhibitors to determine the inhibitors’ effect on cellular entry. Integrins signal bidirectionally via “inside-out” and “outside-in” signaling.^21,36-41^ Inside-out signaling is initiated by intracellular signaling upstream of talin and other adaptor proteins binding to the integrin β-subunit cytoplasmic tail (β-CT), which causes integrin extension (ECS) and concomitant increases in high-affinity ligand binding.^20, 21^ Integrin engagement with macromolecular ligands stimulates the transient exchange of talin for Gα_13_’s occupancy of the β-CT^42, 43^ which initiates integrin outside-in signaling. In the context of viral infection, integrin outside-in signaling induces cell spreading, retraction, and internalization of integrin-associated ligands. We used cell-permeable inhibitors of integrin outside-in and inside-out signaling^42^ to test the role of canonical integrin signaling during cell entry of SARS-CoV-2^R18^ and infectious SARS-CoV-2. Taken together, our results demonstrate that integrins play a significant role in the infectivity of SARS-CoV-2.

## RESULTS

### Integrin extension promotes SARS-CoV-2^R18^ cell entry

To facilitate studies of SARS-CoV-2 host-cell entry outside of BSL-3 containment, we generated UV-inactivated virus particles. Under our experimental conditions, a minimum UV dose of 100 mW•sec/cm^2^ was sufficient to completely inactivate 10^7^ virions/ml distributed in 500 μl samples of a twelve well plate (**Fig. 2A-B**). UV-inactivated virus samples were fluorescently labeled with a lipophilic lipid probe, octadecyl rhodamine B (R18), which intercalates into the envelope membrane.^44^ Labeled samples were purified and characterized as we have described previously for Sin Nombre virus.^45^

**Figure 2.**
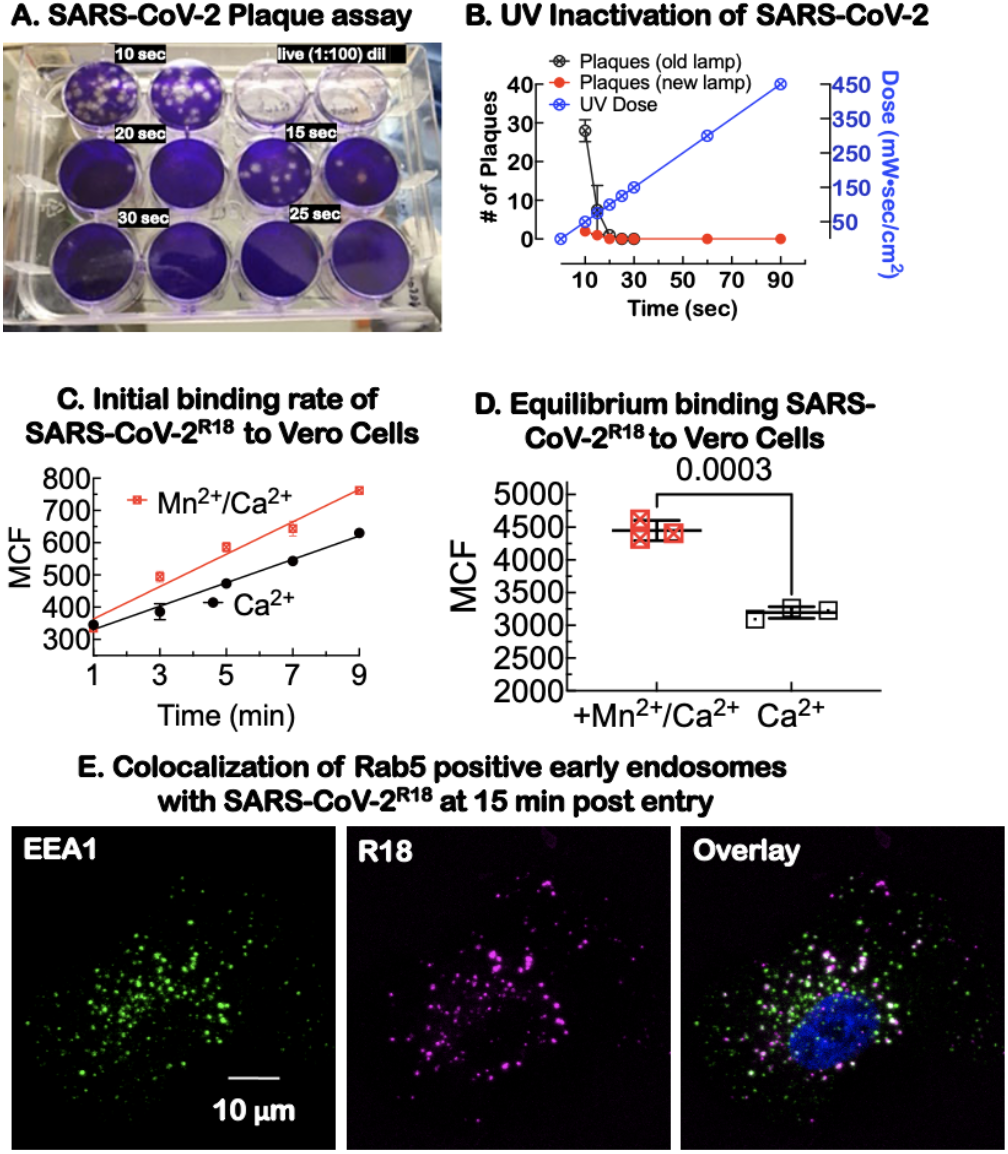
Characterization of UV-inactivated virus for Sars-Cov-2 studies. **A.** Duplicate plaque assays of supernatants of Sars-CoV-2 exposed to increasing doses of 254 nm radiation and then tested for viability. The live virus completely lysed the cells at 1:100 dilution relative to U.V. exposed virions. **B.** Graph shows UV dose response, leading to a significant decrease in plaque forming units at different doses. For our experiments, 90 second (450 mW•sec/cm^2^) U.V. dose was used to inactivate the virus prior to removal from the BSL-3 laboratory. **C.** Binding kinetics of SARS-C0V-2^R18^ to suspension Vero E6 cells at 37°C in Ca^2+^ and Ca^2+^/Mn^2+^ replete cells. **D.** Equilibrium binding of SARS-CoV-2R18 after 30 min incubation at 37°C. **E.** Confocal microscopy imaging of cells after incubation with SARS-CoV-2^R18^ (magenta) for 15 min, then fixed and labeled for early endosome marker, early endosome antigen 1 or EEA1 (green) an effector protein for Rab5, and nuclei (Hoechst 33258, blue). SARS-CoV-2^R18^ vesicles are trafficked to the perinuclear region and a subset are co-localized with EEA1. Images are maximum projections and have been brightness and contrast enhanced.

We hypothesized that activation of integrins by Mn^2+^, ^23, 36^ which induces integrin extension and higher ligand affinity, would provide favorable spatial orientation of the RGD-binding motifs to facilitate SARS-CoV-2^R18^ binding (**Fig. 1A**). Therefore, we measured initial rates (<10 min binding time) of binding in activated cells (Mn^2+^/Ca^2+^) relative to resting (1 mM Ca^2+^ only). The slopes of the graphs were 4.4 and 1.3 for activated and resting samples, respectively, indicating that Mn^2+^ activation accelerated cell entry dynamics **(Fig. 2C)**. However, at equilibrium (>20 min incubation), the cell occupancy of SARS-CoV-2^R18^ was only ~ 20 % higher in Mn^2+^-treated samples **(Fig. 2D)**.

To assess the ability of SARS-CoV-2^R18^ to enter cells, we used confocal microscopy to image the relative distribution of SARS-CoV-2^R18^ and EEA1, an effector protein for Rab5 positive early endosomes. Adherent cells were incubated on a coverslip with SARS-CoV-2R18 for 15 minutes, washed, fixed, and immunolabeled for EEA1 **(Fig. 2E)**. Internalized SARS-CoV-2^R18^ was frequently found in EEA1 positive early endosomes and perinuclear space, demonstrating that the SARS-CoV-2^R18^ internalizes and traffics as one might expect.^29-34^ Together, these results show that UV-inactivated SARS-CoV-2^R18^ is a useful probe for the investigation of SARS-CoV-2 entry mechanisms.

### Inhibition of integrin activation or binding to SARS-CoV-2^R18^ blocks intracellular trafficking

To further investigate the role of integrins in SARS-CoV-2^R18^ entry into Vero E6 cells, we used high binding affinity integrin antagonists: 1) BTT 3033, a selective antagonist (EC_50_ = 130 nM) of integrin α_2_β_1_ that binds to a site close to the α_2_I MIDAS domain and stabilizes the integrin bent conformation state (BCS),^46^ 2) ATN-161, a non-RGD peptide^47^ derived from the synergy region of fibronectin,^48^ known to exhibit specific antagonism for α_5_β_1_ and α_IIb_β_3_ and also recently shown to inhibit SARS-CoV-2 infectivity,^26^ and 3) GLPG0187, a high-affinity, broad-spectrum (EC_50_ <10 nM) integrin receptor antagonist of RGD integrins α_5_β_1_, α_v_β_3_, α_v_β_5_, α_v_β_1_, α_v_β_6_. ^49^ We used a titrated, 5-fold excess of unlabeled SARS-CoV-2 relative to fluorescent SARS-CoV-2^R18^ as a control for competitive inhibition of SARS-CoV-2^R18^ binding.

Paired samples of cell suspensions in Mn^2+^-replete and Mn^2+^-free media were treated with the above integrin antagonists. Total viral binding was normalized to Mn^2+^-treated samples for each experimental condition. Mn^2+^ treatment increased SARS-CoV-2^R18^ occupancy of cells by ~20% compared to Mn^2+^-free conditions (**Fig. 3A-C**). The positive control for inhibition (5xCov-2 in data graphs) blocked 80% of SARS-CoV-2^R18^ and was agnostic to Mn^2+^ treatment. Reasoning that the residual signal of 5x Cov-2 treated samples was due to non-specific binding to the cell membrane, we subtracted the fluorescence of cells blocked with 5xCov-2 and then normalized the data to mock treated cells. We compared the relative efficacy of the inhibitors in Mn^2+^-replete and -free conditions of the normalized data (**Fig. 3D-F**). The fraction of Mn^2+^-activated integrins (20%) were refractory to BTT 3033 treatment (**Fig. 3D**). BTT 3033 selectively binds to the BCS integrin structure^46^ and does not bind to Mn^2+^ activated integrins. In contrast, ATN-161 and GLPG0187 were agnostic to Mn^2+^ treated cells, as the same baseline was achieved for either condition (**Fig. 3D-F**). The inhibition is subtly better in the presence of Mn^2+^ due to higher affinity for the RGD-targeting inhibitors. Overall, GLPG0187 was a better competitive inhibitor of SARS-CoV-2^R18^ compared to ATN-161. The mechanistic specificity of integrin inhibition by these antagonists (BTT 3033 vs. GLPG0187) in regards to SARS-CoV-2 uptake strongly supports the ideas that: 1) integrin RGD engagement is an essential co-factor for cell entry and 2) integrin extension is required for cell entry based on BTT 3033’s mechanism of action.

**Figure 3.**
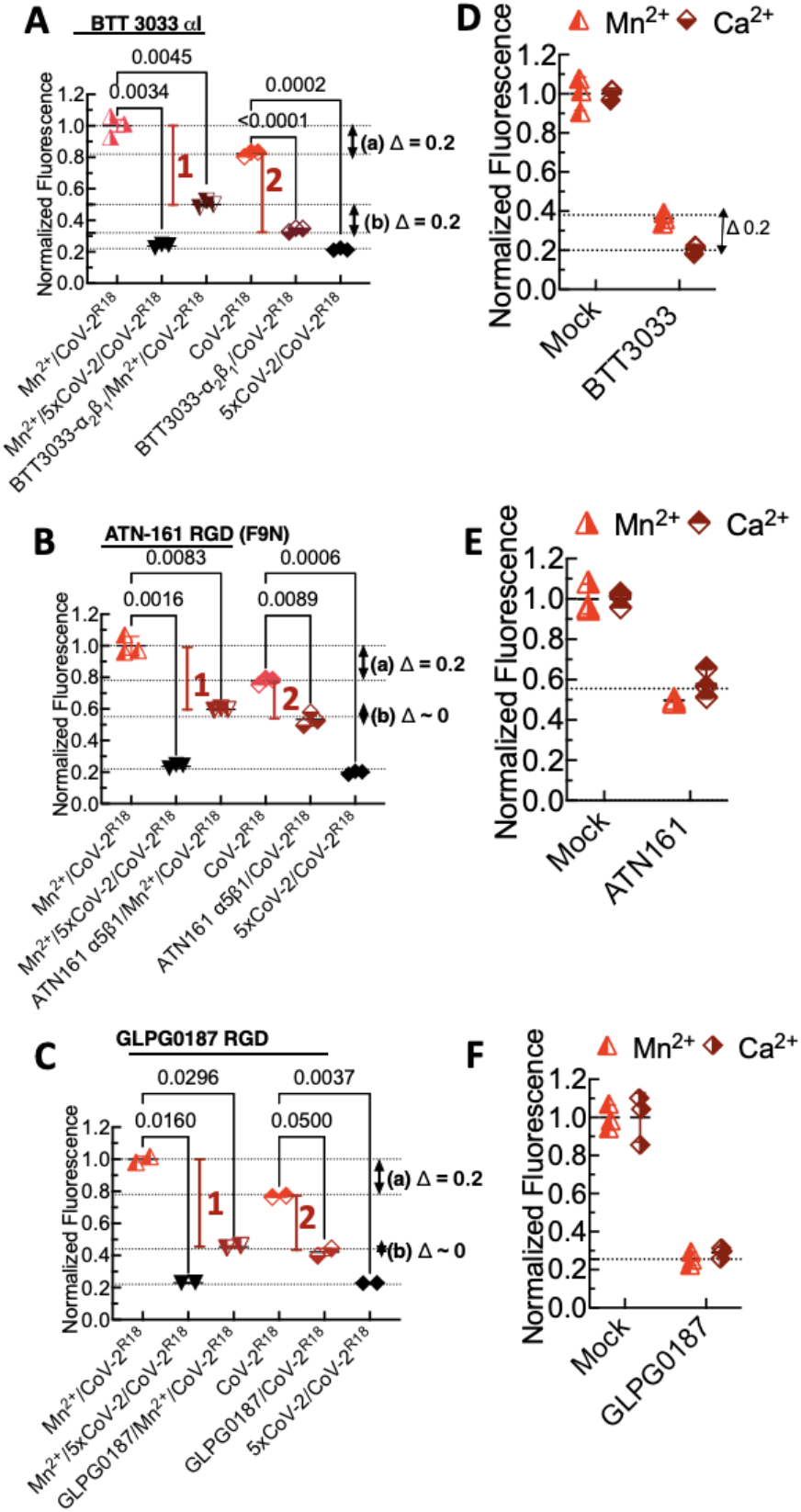
Flow cytometry assays of binding inhibition of SARS-CoV-2^R18^ shows specificity of αI allosteric antagonist BTT 3033, RGD fibronectin synergy domain (F9N) ATN-161, and broad spectrum RGD antagonist, GLPGO187. Vero E6 suspension cells in 40 μl volumes (1,000 cells/μl) were first incubated with 10 μM integrin antagonists or 5x unlabeled Sars-CoV-2 (CoV-2) in ±Mn^2+^ media for 20 min at 37°C. Sars-CoV-2^R18^ was then added (5,000 SARS-CoV-2^R18^/cell) to the tubes and incubated for another 20 min. The samples were centrifuged and resuspended in 95 μl HHB buffer and analyzed on a flow cytometer. **A-C.** Total binding data were normalized to +Mn^2+^ samples. The p-values between samples are indicated on the horizontal bar. **A. BTT 3033.** Red vertical bars 1 and 2 denote difference between mock- and inhibitor-treated samples for Mn^2+^-replete and Mn^2+^-free samples. *(a) Δ = 0.2*, refers to relative difference in fluorescence intensity due to SARS-CoV-2^R18^ in Mn^2+^ replete samples and Mn^2+^ free samples. *(b) Δ = 0.2*, refers to the fractional difference inhibition of SARS-CoV-2^R18^ binding by BTT 3033 in Mn^2+^ replete cells and Mn-free samples. The annotations have similar meanings for **B.** ATN and **C.** GLPG0187. **D-F.** Specific binding of SARS-COV-2^R18^ to cells determined by subtraction of non-specific binding represented by 5xCov-2 data points. The data are normalized to mock treated cells.

We then used live-cell confocal microscopy to visualize Vero E6 cell entry and trafficking of SARS-CoV-2^R18^ in DMSO (mock)- and BTT 3033-treated cells (**Fig. 4A, B**). Most of the cells treated with GLPG0187 de-adhered from the plate and were thus not suitable for imaging. The loss of cells with GLPG0187 was likely due to the loss of integrin-mediated adhesion by the broad-spectrum inhibitor. Cells were imaged at 3-min intervals for 21 min after addition of ~10^7^ SARS-CoV-2^R18^ particles. In DMSO treated cells (Mock in **Fig. 4**), SARS-CoV-2^R18^ particles were visible at cell membranes within 3 minutes subsequently developed punctate features at the cell periphery and trafficked to the perinuclear space. The rate of cell entry (time to perinuclear space ~ 10 min) was comparable to infectious virions.^50^ For the BTT 3033-treated cells, early peripheral membrane localization of SARS-CoV-2^R18^ showed significant diminution of discernable puncta and did not undergo retrograde traffic towards the perinuclear region within the timeframe of the experiment. The relative amount of virus binding to the surface was also reduced with BTT 3033 treatment (**Fig. 4A, B**), consistent with reduced binding observed by flow cytometry measurements (**Fig. 2**).

**Figure 4.**
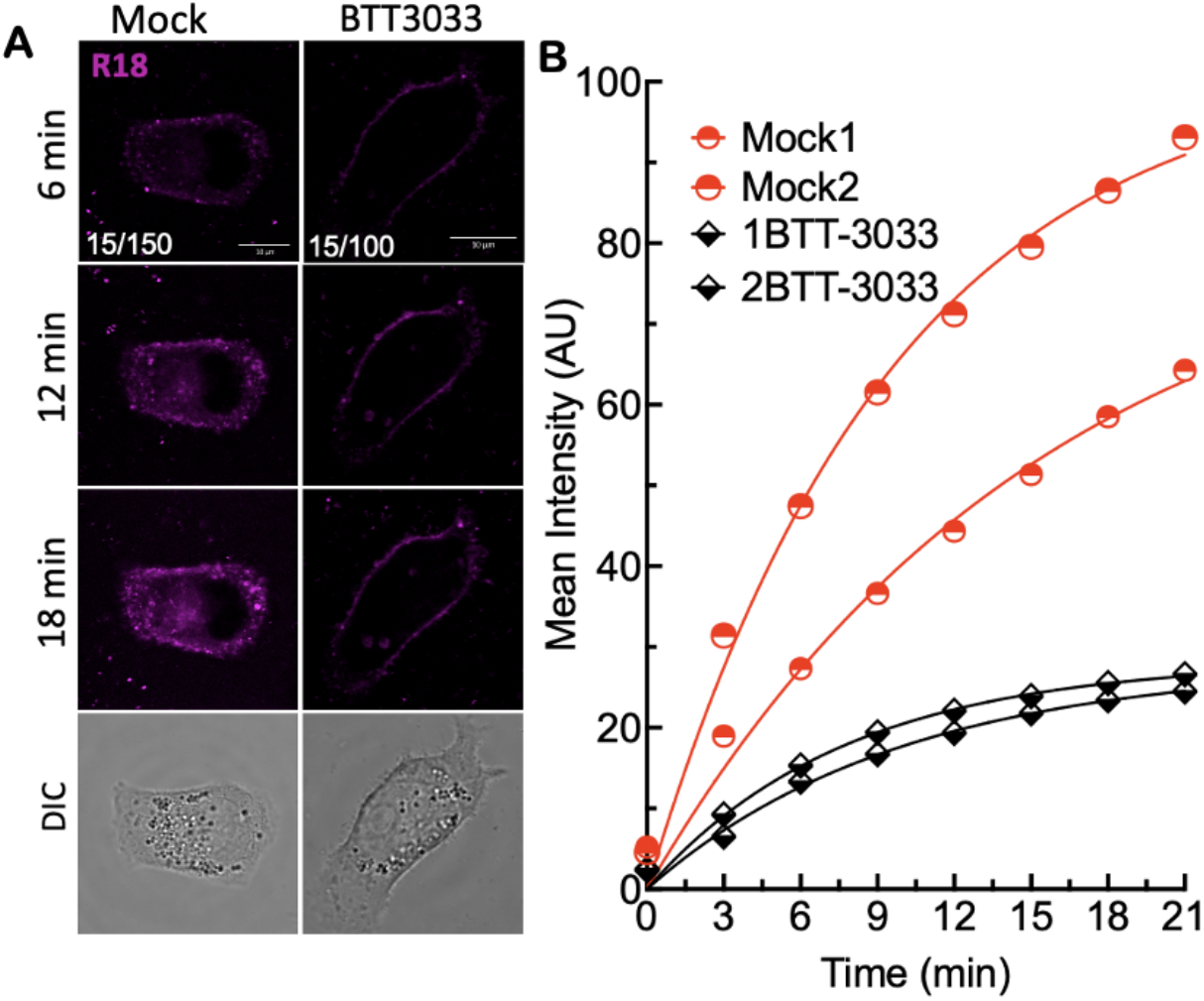
Stabilization of bent closed confirmation with aI MIDAS domain binding integrin antagonist inhibits intracellular trafficking of SARS-CoV-2^R18^. **A.** Live cell imaging of SARS-CoV-2^R18^ (magenta) binding and endocytosis shows perinuclear localization of SARS-CoV-2^R18^ vesicles while virus is seen to accumulate at the plasma membrane when α_2_β_1_ integrins are inhibited by 10 μM BTT 3033. Fluorescence images represent maximum projections of five confocal z slices. Mock and BTT 3033 treated samples are shown with different lookup tables (LUT) since binding in treated cells was lower than untreated cells. LUT lower/upper values are presented in lower left corner of 6 minute timepoint images. Scale bars, 10 μm. **B**. Traces of absolute intensity values of virus binding over time. Two representative cells for each condition are plotted from data acquired on the same day to enable direct comparison of intensity values. Data were fit to a non-linear regression function with arbitrary constants for appearance purposes.

### Inhibition of Gα_13_-integrin β-subunit interaction blocks SARS-CoV-2^R18^ cell entry

Canonical, physiologic integrin activation is started by sequential waves of inside-out signaling initiated by talin binding to the β-subunit cytoplasmic tail (β-CT), which causes integrin extension (ECS in Fig. 1).^51^ ECS integrin binding to immobilized ligand facilitates outside-in signaling as a force applied at the adaptor protein by the actin cytoskeleton is resisted at the ligand-binding site.^20, 21^ Transmission of the tensile force through the integrin to the adaptor protein stabilizes high-affinity integrin binding (in the EOS) to the ECM. This induces the G protein subunit Gα_13_ to transiently replace talin^43^ at the β-CT site, leading to cell spreading, retraction, migration, and internalization of the receptor^40, 52-54^ (**Fig. 5A**). The sequential mechanism of inside-out and outside-in was confirmed in part by the use of two myristoylated peptides; mP6 (Myr-FEEERA-OH), derived from the Gα_13_-binding ExE motif of integrin β-CT, and mP13 (Myr-KFEEERARAKWDT-OH) mimicking the β-CT’s talin binding domain.^42^ mP6 is known to suppress the early phase of outside-in signaling and mP13, which binds both talin and Gα_13_ blocks all phases of integrin signaling.^42^ To investigate the relationship between the integrin signaling events and SARS-CoV-2 engagement and cell entry, we treated cells with mP6 peptide, which inhibited cell entry of SARS-CoV-2^R18^ in flow cytometry and microscopy experiments (**Fig. 5B-D**). Similarly, mP13 inhibited cell entry in flow cytometry experiments (data not shown). The results for mP6 treated cells (**Fig. 5B-D**) suggest that SARS-CoV-2 engagement initiates a Gα_13_-mediated outside-in integrin activation inhibited by mP6, as previously demonstrated for the Sin Nombre virus.^23^

**Figure 5.**
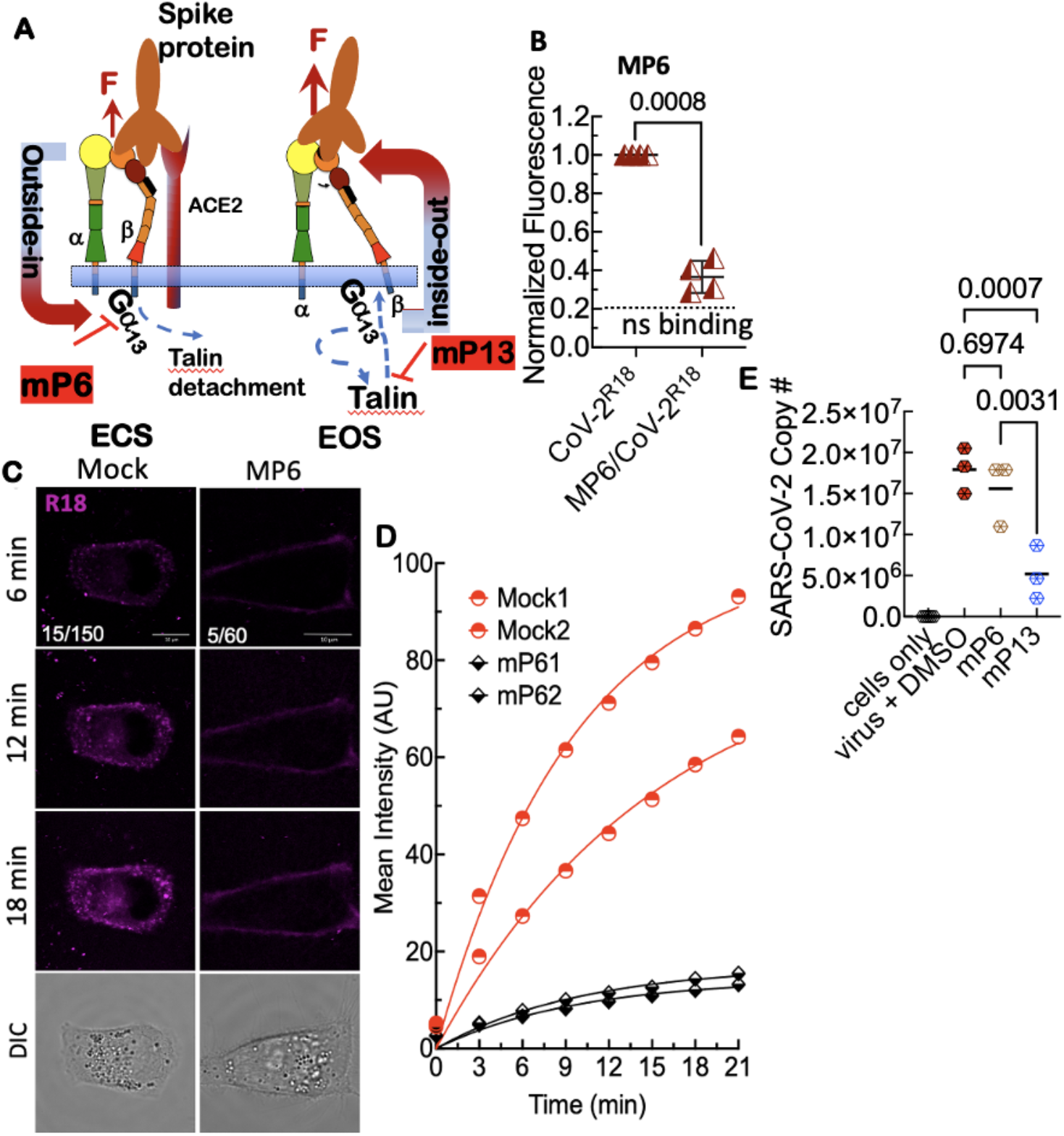
Inhibition of outside-in integrin signaling blocks cell entry of SARS-CoV-2^R18^, suggesting integrin-mediated signaling is required for cellular trafficking. **A.** Model of outsideinside-out signaling for integrin-mediated cell entry. Hypothetical SARS-CoV-2 binding to integrin β_1_ initiates Gα_13_ binding to the β_1_ cytoplasmic tail, which stimulates outside-in signaling in the absence of a known receptor-stimulated GPCR mediated inside-out signaling. mP6 is a specific inhibitor of Gα_13_ binding to the β_1_ cytoplasmic tail. **B.** Relative fluorescence readings of suspension Vero E6 cells after 30 min incubation with SARS-CoV-2^R18^ in vehicle and 100 μM mP6 treated cells. **C**. Live cell imaging of SARS-CoV-2^R18^ (magenta) binding and endocytosis shows cell membrane and perinuclear localization of SARS-CoV-2^R18^ vesicles while virus is seen to remain at the plasma membrane in cells treated with 50 μM mP6. LUT ranges shown in the bottom left corner of 6-min timepoint images. Scale bars, 10 μm. **D**. Traces of absolute intensity values of virus binding over time. Two representative cells for each condition are plotted from data acquired on the same day to enable direct comparison of intensity values. For comparison, mock-treated cell data are the same as in Figure 4. Data were fit to a non-linear regression function with arbitrary constants for appearance purposes. **E.** Inhibition of SARS-CoV-2 productive infection. Suspension Vero E6 cells were preincubated with 250 μM mP6 and 250 μM mP13 for 30 min and followed by infection with 0.01 MOI SARS-COV-2 for an additional 60 min incubation. Cells were washed twice and transferred to a 12 well plate for 48 hours and assayed for viral RNA by RT-qPCR.

### Blocking of integrin signaling significantly inhibits productive infection of cells by SARS-CoV-2

Because mP6 and mP13 are membrane-permeable peptides, they were suitable for infectivity experiments while obviating the need to expose cells to DMSO for extended periods. We therefore tested the efficacy of mP6 and mP13 at inhibiting cell entry and productive infection in Vero E6 cells with a 0.01 multiplicity of infection (MOI) of SARS-CoV-2. For the productive infection assay, infected cells were plated at confluency (500,000 cells/well in a 12 well plate) to minimize cell growth for 48 hrs postinfection. We used RT-qPCR to measure viral RNA in the suspended cells or intact cell monolayers at 48hrs post-infection, respectively. At 48 hrs post-infection, inhibition of productive infection by mP13 was significant relative to mock-treated cells, whereas the effect of mP6 was insignificant (**Fig. 5E**). These results suggest the binding of Gα_13_ to β_3_ integrin induced by SARS-CoV-2 in resting cells (**Fig. 5B**) is a dispensable mechanism of integrin activation under the prevailing conditions of cells perturbed by viral replication.^23^ Stated differently, the primary infection was conducted with quiescent cells, in which Gα_13_-dependent integrin activation was initiated by SARS-CoV-2 engagement. We suggest that in postinfection cells, the Gα_13_-mediated signaling is expendable, in contrast to talin-mediated signaling (see discussion).^23^ Thus, the efficacy of mP13 as an inhibitor of productive infection suggests that integrins play an enduring role in the lifecycle of SARS-CoV-2. ^17, 24, 28^

## DISCUSSION

This study provides mechanistic evidence for the functionality of extracellular ligand-binding domains of integrin β_1_ and cytoplasmic tails of integrins in general,^24, 28^ which offer possible molecular links between ACE2 and integrins. We show that Mn^2+^, which induces integrin extension and high affinity ligand binding, enhances the cell entry of SARS-CoV-2^R18^. This is consistent with the notion that integrin affinity and/or extension is an essential factor for cell entry. In support of integrin-dependent endocytosis as a pathway of SARS-CoV-2^R18^ internalization, we used broad-spectrum RGD antagonists such as GLPG0187, which inhibited cell entry regardless of integrin activation status. Our study also suggested integrin specificity. BTT 3033, an αI allosteric antagonist that binds to the bent closed conformation of integrin β1 and stabilizes it, supports the possibility of integrin-dependent endocytosis of SARS-CoV-2^R18^ upon receptor binding. In a different framework, our data also show that SARS-CoV-2^R18^ can bind to low affinity and presumptively bent-conformation integrins,^22^ however, in BTT 3033 treated cells, cell entry by SARS-CoV-2^R18^ is inhibited because integrin activation post-SARS-CoV-2^R18^ engagement is prevented. Thus, our data contextualize integrin extension as the “*sine qua non* of integrin cell adhesion function,”^22^ which in turn is an essential condition for integrin-mediated cell entry by SARS-CoV-2.

The binding of adaptor proteins such as talin and Gα_13_ to the integrin β-subunit cytoplasmic tail are essential elements of inside-out and outside-in physiologic integrin signaling. ^42^ mP6 and mP13 both blocked initial cell entry of SARS-CoV2^18^. However, only mP13 inhibited productive infection. In an earlier study of Sin Nombre virus infectivity, we showed that artificially-induced discharge of intracellular Ca^2+^stores elicited integrin inside-out signaling function, which dispensed with the need for Gα_13_ mediated signaling.^23^ Under our experimental conditions lasting 48 hours, we estimate that successive replication and release of progeny virions occurred every ≥8 hours. ^50, 55^ We suggest that the overall process of productive infection, is characterized by viral perturbation of Ca^2+^ homeostasis within the infected cells.^56^ Thus one might expect persistent integrin priming^57, 58^ which may render the Gα_13_ pathway to be redundant.

Mészáros *et al*. ^24^ have used bioinformatics to predict the existence of short amino acid sequences (~3-10 residues): short linear motifs (SLiMs), in the cytoplasmic tails of ACE2 and integrins that mediate endocytosis and autophagy. Some of their theoretical predictions have been validated by experimental studies. First, Kliche *et al*. ^28^ confirmed the existence of SLiMs. They extended their findings to establish a potential connection between ACE2 and integrin β_3_ cytoplasmic tail interactions with scaffolding and adaptor proteins linked to endocytosis and autophagy. Second, SLiM sequences known to bind and activate the transmembrane glycoprotein neuropilin 1 (NRP1) were identified as potential mediators of SARS-CoV-2 endocytosis.^24^ Interestingly, NRP1, which is abundantly expressed in the olfactory epithelium is now declared as an effector for SARS-CoV-2 infection.^34, 35^ NRP1 localizes at adhesion sites and promotes fibronectin-bound, activated α_5_β_1_ integrin endocytosis and directs the cargo to the perinuclear cytoplasm.^29-34^ Studies have shown that the endocytosis of active and inactive integrins to EEA1-containing early endosomes follows distinct mechanisms involving different adaptor proteins. The inactive integrin is promptly recycled back to the plasma membrane via an ARF6- and EEA1-positive compartment in a Rab4-dependent manner. ^31^ We observed that in BTT 3033-treated cells replete with inactive β_1_ integrins, SARS-CoV-2^R18^ remained membrane-bound, whereas untreated cells displayed internalization and perinuclear localization of SARS-CoV-2^R18^. This is consistent with the known trafficking of active integrins, including those directed by NRP1, to the perinuclear space.^29, 30, 32^

Although several integrin types^24-28^ are believed to be co-receptors of SARS-CoV-2 infectivity, our study suggests inhibitor specificity for integrin β_1_. This is consistent with known factors: 1) correlated increased expressions of β_1_ ^14^ and ACE2 in relevant tissues,^15, 16^ 2) cytoplasmic tail *in cis* interactions between ACE2 and integrin β_1_, ^13^ and 3) synergy between ACE2 and integrin β_1_ signaling that promotes RGD mediated cell adhesion.^17^ To optimize integrin engagement, our cell-binding assays and primary infection assays were carried out in suspension such that ACE2 and integrins were not segregated by cell polarization. ^59, 60^ However, our microcopy studies on adherent cells were in agreement with the flow cytometry results. Thus, our study represents an initial step forward in establishing a mechanistic role for SARS-CoV-2-mediated integrin activation required for cell entry and productive infection.

## MATERIALS AND METHODS

### Materials

USA-WA1/2020 SARS-CoV-2 strain was obtained from BEI Resources (NIAID, NIH). Integrin inhibitors, BTT3033, a selective inhibitor of α_2_β_1_, ATN-161 an integrin α_5_β_1_ antagonist,^26^ and GLPG0187 a broad-spectrum integrin inhibitor, were purchased as powders from Tocris Bioscience. The EEA1 rabbit monoclonal antibody (clone C45B10) was from Cell Signaling Technologies (CAT# 3288S). Alexa fluor 647 conjugated F(ab’)2 fragment goat anti-rabbit IgG was from Invitrogen (CAT# A21246). In addition, myristoylated peptides; mP6 (Myr-FEEERA-OH) and mP13 (Myr-KFEEERARAKWDT-OH) were custom synthesized at Vivitide.

### Cell culture

African green monkey kidney cells (Vero E6, ATCC) were maintained in DMEM media from Sigma CAT# D5796. All media contained 10% heat-inactivated fetal bovine serum (FBS), 100 U/ml penicillin, 100 μg/ml streptomycin, and 2 mM l-glutamine and were kept at 37 °C in a CO_2_ water-jacketed incubator of 5% CO_2_ and 95% air (Forma Scientific, Marietta, OH, USA).

### UV inactivation and fluorescent labeling of the envelope membrane of SARS-CoV-2 with octadecyl rhodamine (R18)

USA-WA1/2020 SARS-CoV-2 strain (from BEI Resources, NIAID, NIH) was cultured in Vero E6 cells in a biosafety level 3 (BSL-3) containment under a protocol approved by the University of New Mexico’s Institutional Biosafety Committee or IBC (Public Health Service registration number C20041018-0267). Live SARS-COV-2 were harvested at peak titers of 10^7^ plaque-forming units/mL (PFU/ml). Next, SARS-CoV-2 was U.V. inactivated using 254 nm (≈ 5 mW/cm^2^) U.V. irradiation of a TS-254R Spectroline UV Transilluminator (Spectronics Corp., Westbury, NY) following a similar protocol for inactivating pathogenic orthohantaviruses. ^45, 61^ Briefly, Vero E6 cells were inoculated with SARS-CoV-2 and maintained at 37°C for 2-4 days. At 70-75% cell death (due to viral cytopathic effect), the supernatant was harvested and subjected to light centrifugation (1000 rpm, 10 min) to remove cellular debris. For U.V. inactivation, supernatants were added to a 12 well plate at 500 μl aliquot/well. Then U.V. irradiated at 3.8 cm above the sample for 0, 10,15, 20, 25, 30, 60, and 90 seconds and then tested for viability by a three-day plaque assay as described elsewhere. ^62, 63^ The titration of U.V. irradiation times was used to establish a minimal U.V. dose for complete inactivation. After U.V. treatment, the 500 μl fractions were pooled into 15 mL tubes stored in a −80°C freezer pending the results of a plaque assay. Under our experimental conditions, we established that a minimum U.V. irradiation interval of 25 seconds was required for the complete inactivation of SARS-CoV-2. A 90 sec UV dose was approved by the IBC for removal of inactivated SARS-CoV-2 out of the BSL-3 lab after it was established that the virus particles were capable of specific binding to Vero E6 cells.

Crude UV-inactivated SARS-CoV-2 samples were purified by floating 10 ml of Vero E6 SARS-CoV-2 supernatant on a density gradient comprising 2 ml volumes of 1.2 g/ml and 1.0 g/ml CsCl in PBS media in 14 × 89-mm Beckman polyallomer tubes. The samples were centrifuged for 1.5 hours at 4 °C using a Beckman SW41Ti rotor at 30,000 pm. Individual fractions were collected, and a refractometer was used to identify the fraction that contained SNV by its putative density of 1.0 g/ml. The purified SARS-CoV-2 samples were stored in 1.0 ml aliquots at −80 °C. SARS-CoV-2 particles were fluorescently labeled and calibrated according to the same protocol used for the Sin Nombre virus (SNV). ^45^ SARS-CoV-2^R18^ particles were stored at −80°C in 20 μl (2×10^8^ particles/μl) aliquots.

### Flow Cytometry Binding Assays of SARS-CoV-2^R18^ to Vero E6 cells

For flow cytometry assays, cells were cultured in T25 or T75 flasks to 80% confluence. Cells were then treated with 0.25% trypsin and then transferred to minimum essential medium (MEM) media. Test suspension cell samples were transferred to microfuge tubes in 40 μl-aliquots (1,000 cells/μl). SARS-CoV-2^R18^ was added to tubes at 5,000 SARS-CoV-2^R18^/cell and incubated using a shaker at 500 rpm for 20 min at 37°C. For blocking assays, cells were incubated with 5x unlabeled SARS-CoV-2 or 10 μM integrin inhibitors for 20 min before the addition of SARS-CoV-2^R18^. Samples were centrifuged at 3,000 rpm; the pellet was resuspended in HHB buffer (30 mM HEPES, 110 mM NaCl, 10 mM KCl, 1 mM MgCl_2_•6H_2_O, and 10 mM glucose, pH 7.4) buffer and read on an Accuri flow cytometer. For kinetic assays, Vero E6 suspension cells in 40 μl volumes (1,000 cells/μl) were placed in ±Mn^2+^ media in duplicate microfuge tubes at 37°C. Sars-CoV-2^R18^ was then added (5,000 virions/cell) to the tubes and incubated for 1, 3, 5, 7, 9 min. At each time point, the tubes were quenched in an ice bath, then samples were centrifuged and resuspended in 95 μl HHB buffer and analyzed on a flow cytometer

### Live Cell Confocal Microscopy

Imaging was performed using a Leica TCS SP8 Laser Scanning Confocal Microscope with a 63× water objective and a Bioptechs objective heater to maintain cells at physiological temperature (~36-37 °C). Vero E6 cells were plated in eight-well Lab-Tek (Nunc) chambers at a density of 30,000 cells per well 24 hours before imaging. Cells were imaged in Tyrode’s buffer (135 mM NaCl, 10 mM KCl, 0.4 mM MgCl2, 1 mM CaCl_2_ 20 mM glucose, 0.1% BSA, 10 mM HEPES, pH 7.2). For integrin inhibition, cells were treated with 10 μM BTT3033-α_2_β_1_ or 50 μM MP6 in Tyrode’s buffer for 30 minutes before imaging. ~1×10^9^ SARS-CoV-2^R18^ particles were added per well and z-stacks (300 nm thickness) were acquired every 3 minutes for 21 minutes to visualize viral cell entry. R18 was excited using 561 nm light, isolated from the white light source. R18 emission and differential interference contrast (DIC) transmitted light were captured with Leica Hybrid detectors (HyD) in a spectral window of 571-636 nm (for R18 emission). Analysis of the accumulation of SARS-CoV-2^R18^ particles in Vero E6 cells was completed using Matlab. Briefly, regions of interest (ROI) were created around the cell membrane and the mean SARS-CoV-2^R18^ intensity was measured within the cell mask at each time point.

### Immunofluorescence

Vero E6 cells were plated on 18-mm coverslips overnight in a 6 well plate at a density of 100,000 cells/well. Cells were exposed to ~1×10^9^ SARS-CoV-2^R18^ particles/well for 15 min at 37 °C, in the presence or absence of 10 μM BTT 3033. Cells were then washed in phosphate-buffered saline (PBS) and fixed using 4% paraformaldehyde (PFA) in PBS for 15 minutes at room temperature. Cells were extensively washed with 10 mM Tris (pH 7.4) and PBS and permeabilized with 0.1% Triton. Cells were labeled with anti-EEA1 primary antibody and anti-rabbit Alexa Fluor 647 secondary. Nuclei were stained with Hoechst 33258. Cells were mounted on microscope slides using Prolong Diamond Antifade Mountant (Invitrogen, CAT#P33970). Samples were imaged using a TCS SP8 Laser Scanning Confocal Microscope with a 63× oil objective.

### Infection Inhibition

Vero E6 cells grown at 80% confluency were trypsinized and divided into microfuge tubes aliquots of 1.5×10^6^cells in 750 μl media containing 250μM mP6, 250μM mP13, 10μM BTT 3033, DMSO, and media only. Samples were shaken at 500 rpm for 30 min at 37°C. After transfer to a BSL-3 laboratory, 0.01 MOI of SARS CoV-2 (lot #P3: 1.2×10^7^ pfu) and then incubated for 60 min while shaking. Tubes were spun down (1000rpm for 3 min), resuspended in fresh media, and spun down again. The cells were then resuspended in 300 μl of media and transferred to a 12 well plate in 100 μl aliquots (500,000 virions/well) for triplicate measurements. An additional 400μl were added to each well for a final volume of 500 μl. The plate was transferred to an incubator for 48 hrs to allow the virus to replicate. The cells were then washed with 1xPBS before RNA was extracted with TRIzol™ (Thermofisher, #15596026) according to the manufacturer’s protocol: (https://assets.thermofisher.com/TFS-Assets/LSG/manuals/trizol_reagent.pdf).

The RNA was quantified with a Nanodrop and cDNA transcribed with the Applied Biosystems™ High-Capacity cDNA Reverse Transcription Kit (Fisher Scientific #43-688-14).

RT-qPCR was performed with TaqMan Fast Advance Kit (Fisher Scientific #4444963) and the following primers ordered from Integrated DNA Technologies: Forward: 5’-CCCTGTGGGTTTTACACTTAA-3’, Reverse: 5’-ACGATTGTGCATCAGCTGA-3’, and probe: 5’-[FAM] CCGTCTGCGGTATGTGGAAAGGTTATGG [BHQ1]-3’ RT-qPCR was analyzed using an Applied Biosystems QuantStudio 5 instrument.

## Acknowledgements

This project is supported by an award from the National Center for Advancing Translational Sciences, National Institutes of Health under grant number UL1TR001449, NIH R35GM-126934 (DSL), Fluorescence microscopy was performed in the University of New Mexico Comprehensive Cancer Center fluorescence microscopy shared resource (NIH P30CA118100).

